# Optimal homotopy analysis of a chaotic HIV-1 model incorporating AIDS-related cancer cells

**DOI:** 10.1101/097865

**Authors:** Jorge Duarte, Cristina Januário, Nuno Martins, C. Correia Ramos, Carla Rodrigues, Josep Sardanyès

## Abstract

The studies of nonlinear models in epidemiology have generated a deep interest in gaining insight into the mechanisms that underlie AIDS-related cancers, providing us with a better understanding of cancer immunity and viral oncogenesis. In this article, we analyse an HIV-1 model incorporating the relations between three dynamical variables: cancer cells, healthy *CD*4+ T lymphocytes and infected *CD*4+ T lymphocytes. Recent theoretical investigations indicate that these cells interactions lead to different dynamical outcomes, for instance to periodic or chaotic behavior. Firstly, we analytically prove the boundedness of the trajectories in the system’s attractor. The complexity of the coupling between the dynamical variables is quantified using observability indices. Our calculations reveal that the highest observable variable is the population of cancer cells, thus indicating that these cells could be monitored in future experiments in order to obtain time series for attractor’s reconstruction. We identify different dynamical behaviors of the system varying two biologically meaningful parameters: *r*_1_, representing the uncontrolled proliferation rate of cancer cells, and *k*_1_, denoting the immune system’s killing rate of cancer cells. The maximum Lyapunov exponent is computed to identify the chaotic regimes. Considering very recent developments in the literature related to the homotopy analysis method (HAM), we construct the explicit series solution of the cancer model and focus our analysis on the dynamical variable with the highest observability index. An optimal homotopy analysis approach is used to improve the computational efficiency of HAM by means of appropriate values for the convergence control parameter, which greatly accelerate the convergence of the series solution.

## 1. Introduction

Nowadays, over 60 million people worldwide have been infected with human immunodeciency virus (HIV), more than 80% of whom live in developing countries. For HIV-infected individuals, cancer remains a significant burden. In particular, the *Kaposi’s sarcoma* (KS) is the most common neoplasm that occurs in patients with AIDS (AIDS-KS). KS is a cancer that develops from the cells that line lymph or blood vessels. It usually appears as tumors on the skin or on mucosal surfaces such as inside the mouth, but tumors can also develop in other parts of the body, such as in the lymph nodes, the lungs, or digestive tract [1]. The epidemic AIDS-KS is the most common type of KS in the United States. This type of KS develops in people who are infected with HIV, the virus that causes AIDS, but KS is not caused by HIV but by a herpesvirus. A person infected with HIV (that is, who is HIV-positive) does not necessarily have AIDS. The virus can be present in the body for a long time, often many years, before causing major illness. The disease known as AIDS begins when the virus has seriously damaged the immune system, which results in certain types of infections or other medical complications, including KS. When HIV damages the immune system, people who also are infected with a certain virus (e.g., the Kaposi sarcoma associated herpesvirus) are more likely to develop KS [2].

Gaining insight into the epidemiology and mechanisms that underlie AIDS-related cancers can provide us with a better understanding of cancer immunity and viral oncogenesis. How can the combination of immunosuppression and activation of inflammation promote cancer development? Our purpose in this paper is to try to give a glancing analysis using a simple dynamical model.

The use of mathematical models as an aid in understanding features of virus infection dynamics (such as HIV-1) has been substancial in the past 20 years. There are two ways for HIV-1 to disseminate i*in vivo* [3, 4, 5]: (i) circulating free viral particules to T cells directly, or (ii) through HIV-infected T cells to healthy T cells (see also [6, 7, 8]). Most of these models focus on cell-free virus spread in the bloodstream [9, 10]. A model concerning the cell-to-cell spread of HIV-1 is relevant, since understanding the dynamics of the HIV infection within lymphatic tissues is vital to uncovering information regarding cellular infection and viral production [11]. In this sense, recent experimental works have provided new insights into the mechanisms underlying HIV-1 cell-to-cell infection and their impact on drug therapy [12].

The model studied here appeared in [13] as a dynamical system accounting for the cell-to-cell spread of HIV-1 together with cancer cells in tissue cultures. This model is aimed at explaining some quantitative features concerning cancer occuring during HIV-1 infection that are unusual and, in the absence of a model, perplexing. The basic starting point of this model has three parts. First, the cancer cells are caused by the changes of the normal cells in the individual due to some physical, chemical or biological factor (for instance, a virus such as human papilloma virus (HPV)) - under normal conditions, the healthy cells in our body can mutate into cancer cells with probability of 10^−6^. Second, the cancer cells have anomalies in growth-related genes (i.e., oncogenes and tumor-suppressor genes) that allow them to proliferate faster than normal cells. Third, the immune system can recognize the difference between cancer cells and normal cells, so it can survey them and then carry out its killing function. Here we must notice that cancer cells can also avoid immune system recognition by means of the so-called immunoediting. However, the model we are investigating is not considering this process.

As pointed out in [13], the studied HIV-1 model has a number of steady states whose existence and stability properties are quite consistent with their biological meanings. Periodic solutions and chaos appear alternately along with the changing of the bifurcation parameters. With this HIV-1 model it is possible to investigate the cancer situation in an individual who is infected by HIV-1.

Given the importance of this type of models in the literature, a great deal of numerical algorithms for approximating solutions can be used in diverse computational studies of the nonlinear dynamics. Wthout doubt the numerical algorithms have been extremely important in the study of complex dynamical systems. However, they allow us to analyse the dynamics at discrete points only, thereby making it impossible to obtain continuous solutions. As a consequence, it turns out to be extremely valuable to develop an analytic approximation methodology which should have three fundamental characteristics: (i) it is independent of any small/large physical parameters; (ii) it gives us freedom and flexibility to choose equation-type and solution expression of high-order approximation series; and (iii) it provides us a convenient way to guarantee the convergence of approximation series, using an auxiliary convergence-control parameter.

One such general analytic technique, which has the three advantages mentioned above, used to get convergent series solutions of strongly nonlinear problems is the so-called Homotopy Analysis Method (HAM), developed by Liao [14, 15, 16], with contributions of other researchers in theory and applications.

Frequently, in order to have an effective analytical approach of strongly nonlinear equations for higher values of time *t*, the simple idea is to apply the HAM in a sequence of subintervals of time with a certain step, giving rise to the so-called Step Homotopy Analysis Method (SHAM). In fact, the homotopy analysis methodology is more general in theory and widely valid in practice for the study of nonlinear problems than other analytic approximation procedures. Indeed, this methodology has been successfully applied to solve a wide variety of nonlinear problems (please see for instance [17] - [20] and references therein), particularly, there has been a growing interest in applying HAM to biological models (please see some illustrative examples in [21]).

The paper is organized as follows. We give in *Section 2* a brief description of the HIV-1 model presented in [13]. In *Section 3*, we analytically show that the chaotic attractor is positively invariant. In *Section 4*, we compute the coupling complexity by means of observability indices, which allow us to rank the dynamical variables from more to less observable. An analytical study, using the homotopy analysis methodology, is carried out in *Section 5*. This section contains the explicit series solution (Subsection 5.1) and an optimal homotopy analysis approach of solutions to improve the computational efficiency of HAM (Subsection 5.2). In particular, we obtain for each dynamical variable an optimal value of the HAM converfogence-control h using an appropriate ratio and using the exact squared residual error. Finally, *Section 6* is devoted to significative conclusions.

## 2. The HIV-1 cancer model

The model that we investigate describes the cell-to-cell dynamics between healthy immune cells and HIV-1 infected immune cells, adding another layer of complexity where healthy immune cells attack and kill cancer cells [13]. Cell-to-cell spread of HIV-1 has been widely studied with theoretical and computational models [6, 22, 23, 24] in order to understand several features of *in vitro* experiments done for this type of virus (as well as that of others). The present model is aimed to describe the dynamics of HIV-1 infection in patients with cancer using a mean-field approach. The model is given by the following set of ordinary differential equations:

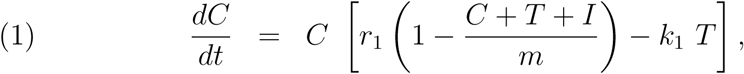

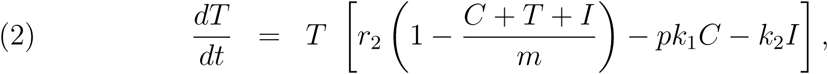

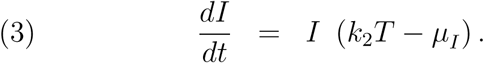

The state variables are the population numbers of cancer cells (*C*), healthy cells (*T*), and HIV-infected cells (*I*). The constants *r*_1_ and *r*_2_ are the uncontrolled proliferation rate of cancer cells and the intrinsic growth rate of healthy cells, respectively, with *r*_1_ > *r*_2_. Notice that the proliferation of cancer cells needs to be higher than the replication of healthy cells, since a common characteristic of cancer cells is their increase in proliferation rate due to genetic anomalies in both oncogenes and tumor suppressor genes [25]. *k*_1_ corresponds to the immune system’s killing rate of cancer cells; *k*_2_ is the infection rate coefficient. Moreover, m is the effective carrying capacity of the system; *p* is the losing rate of the immune cells because of the killing of cancer cells. Finally, the constant *μ_I_* represents the whole immune system killing’s effect on the infected cells.

Notice that the growth of the populations for the cancer and the immune system cells is limited by a logistic-like function, given by 1 — (*C* + *T* + *I*)/*m*. This function introduces competition between the cancer and both the healthy and infected T cells. Competition amongst cells is crucial in tumor dynamics since cancer cells often have a selective advantage in terms of increased proliferation rates (i.e., *r*_1_ > *r*_2_ in the model). The logistic term is not found in Eq. (3) because the population of infected CD4+ T cells arises from the population of healthy CD4+ T cells (variable *T*(*t*)) upon infection.

We will consider throughout our study the same parameter values used in Ref. [13]. Such values were obtained from the literature of clinical and mathematical models. Specifically, we will set *r*_2_ = 0.03, *k*_2_ = 0.0005, *m* = 1500, *p* = 0.1, *μ_I_* = 0.3 and take the uncontrolled proliferation rate of the cancer cell *r*_1_ and the immune system’s killing rate of cancer cells *k*_1_ as control parameters (0.1775 ≤ *r*_1_ ≤ 0.18425 and 0.0001 ≤ *k*_1_ ≤ 0.000107). According to the literature [5, 26, 27], the probability that a healthy cell will become a cancer cell is very small, even if there some factors that urge the transformation. We assume that the cancer is caused by just one cell because of gene mutation.

## 3. Positively invariant sets

In the following Sections we will examine the long-term behavior of the three-dimensional chaotic attractors arising in the HIV-1 system modeled by Eqs. (1)-(3). The chaotic attractor, displayed in Fig. 1, governs the population dynamics of the cancer cells, healthy *CD*4+ T lymphocytes and infected *CD*4+ T lymphocytes, given by C, T and I, respectively. Figure 1(a) displays the chaotic attractor using two different but close initial conditions. The same attractor is plotted in Fig. 1(b) together with the fixed points found in the phase space. Such points are all unstable (see [13] for a full description of the equilibria and stability of the system).

**FIGURE 1.**
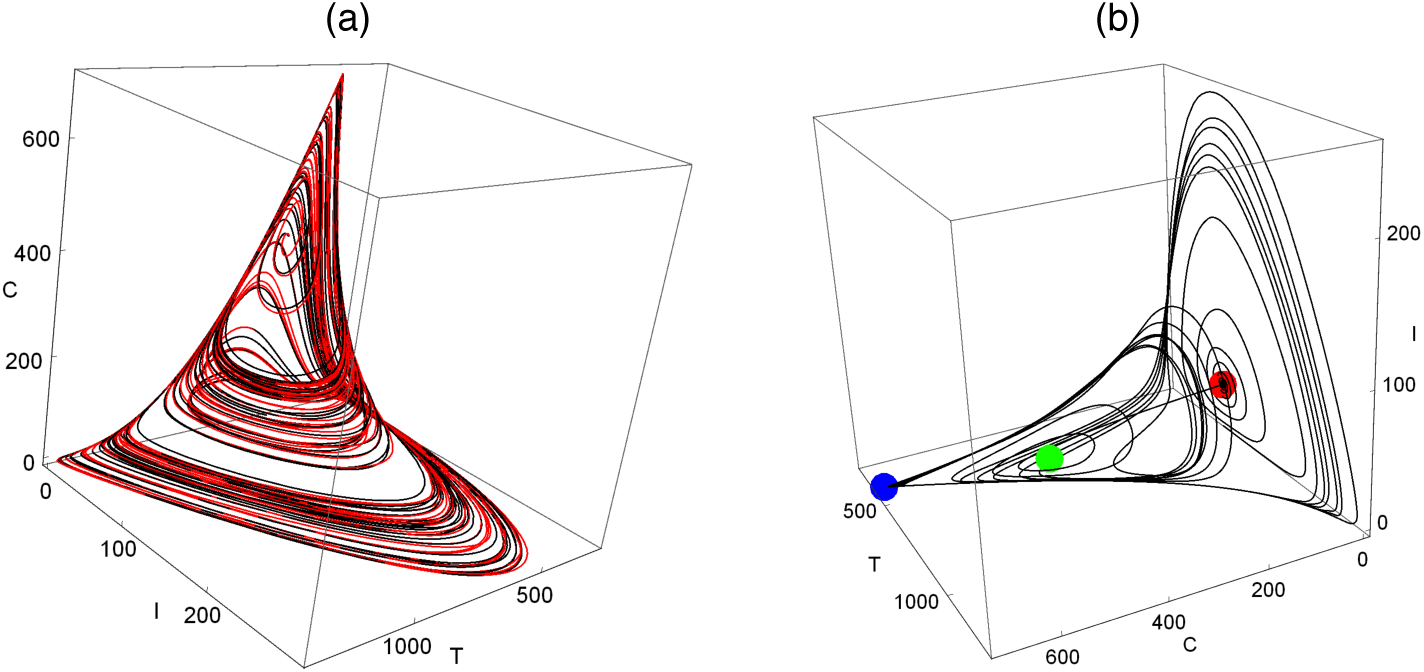
Chaotic attractor corresponding to the HIV-1 system of (Eqs. (1)-((3), for *r*_1_ = 0.1842 and *k*_1_ = 0.0001. (a) Attractor obtained using two different but close initial conditions displayed in black and red. (b) Trajectory converging to the chaotic attractor represented together with the fixed points *E*_4_ (red), *E*\* (green), and *E*_3_ (blue). The names of the fixed points follow the notation in Ref.[13]. Here the fixed point *E*_4_ has complex eigenvalues with real part (—, —, +). The fixed point *E*\* also has complex eigenevalues with real part (—, +, +), and the fixed point *E*_3_ has real eigenvalues with sign (—, —, +).

A preliminary question to answer, before doing any further analysis, is to find conditions for which trajectories will not “escape to infinity”, so that they will remain confined to a compact set. In biological terms, this boundedness means that no population grows without limit and thus the model captures correctly the dynamics (see [28] as an example of boundedness concept application).

Let us consider a new function Ψ = *C* + *T* + *I*, i.e., the sum of the populations involved in the 3D system. The temporal derivative of Ψ is

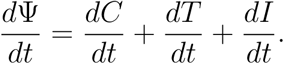

Adding *ε*Ψ to 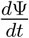, we consider 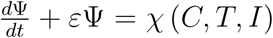, for some *ε* > 0. An upper bound of *χ* (*C*, *T*, *I*) is given by

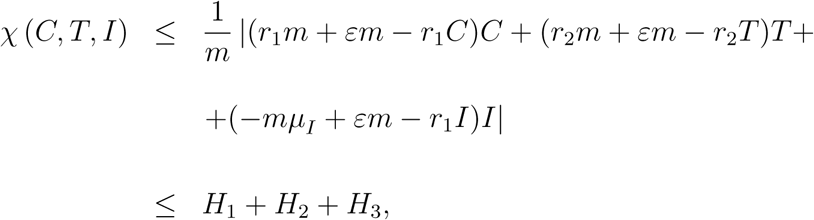

with 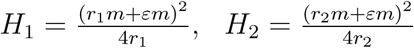,and 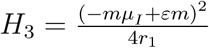.

We obtain *χ* (*C T*, *I*) ≤ *H*_1_ + *H*_2_ + *H*_3_ = *H*. It follows that *d*Ψ/*dt* ≤ —*ε*Ψ + *H*.

Using the differential form of the Gronwall’s inequality [29], we find

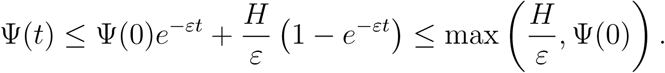

As a consequence, the trajectories starting from any arbitrary initial condition will remain confined to a compact set.

## 4. Observability analysis

When a dynamical system is investigated, there are usually some variables that provide a better representation of the underlying dynamics. More precisely, in a number of practical situations, the choice of the *observable* does influence our ability to extract dynamical information of a given attractor. This fact results, in a considerable degree, from the complexity of the coupling between the dynamical variables. With the computation of *observability indices*, this coupling complexity can be estimated and the variables can be ranked [30, 31, 32]. In the context of nonlinear dynamics, the choice of the observable has a direct relation with problems such as control, model building and synchronization (please see [31] and references therein).

It is important to notice that, despite the potential practical importance of this concept, observability has not been commonly addressed by the research community of nonlinear dynamics. The following method thus provides an illustration of how our understanding of nonlinear problems can be enhanced by the theory of observability. In the next lines, we perform an observability analysis of the three-variable HIV-1 model, which involves a mathematical structure provided by the theory of observability - the definition of the *observability matrix* [30, 31, 32].

Let us consider a dynamical system

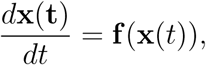

where *t* is the time, x ∈ ℝ^m^ is the state vector and f is the nonlinear vector field. This system is called the *original system*. The observable variable is obtained using a measurement function h : ℝ*^m^* → ℝ, such that *s*(*t*) = *h*(x(*t*)). A system of three ordinary differential equations (*m* = 3) of the form

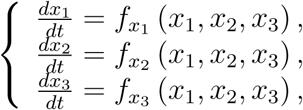

can be reconstructed in a three-dimensional space. More precisely, using the variable *s*, the reconstructed portrait is spanned by the derivative coordinates according to

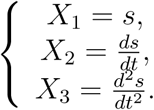

The successive temporal derivatives of *s* constitute a *derivative vector*. The dynamics of this space defined by the three derivative coordinates is expected to be equivalent, in a certain sense, to the dynamics of the system defined by the original coordinates. In order to analyse the quality of the reconstructed space, we study the properties of a coordinate transformation Φ*_s_* between the original dynamical variables and the derivative coordinates,

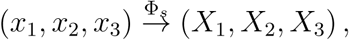

where the subscript *s* indicates the dynamical variable from which the reconstruction is undertaken. For the observable variable *s*, the transformation Φ*_s_* reads

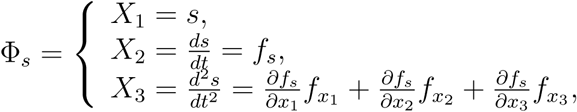

where s can either represent *x*_1_, *x*_2_, or *x*_3_, which are the three components of the vector field f. The coordinate transformation contains information on the nature of the coupling between the dynamical variables “seen from one observable point of view”. For our model we are going to consider three coordinate transformations Φ*x*_1_, Φ*x*_2_ and Φ*x*_3_. In the context of the observability theory, it is critical to investigate in what conditions a dynamical state can be constructed from a single variable and how the nature of the couplings may effect the observability of a given system.

Theoretically, in order to reconstruct a dynamical state from *s*, the striking case occurs when the transformation Φ *_s_* defines a diffeomorphism, i. e.,Φ *_s_* is a continuous invertible function whose inverse is differentiable. In other words, the coordinate transformation Φ *_s_* defines a diffeomorphism from the original phase in the reconstructed one if the determinant of its jacobian matrix, *J* (Φ *_s_*), never vanishes for each point of the phase space.

Thus, the study of the jacobian matrix *J* (Φ *_s_*) is critical and gives us relevant information for the characterization of the coordinate transformation Φ_5_. In particular, the map Φ *_s_* is locally invertible at a given point *x*_0_ if the Jacobian matrix has full rank, i.e., if *rank* 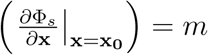. As a consequence, the original dynamical system is locally observable if the previous sufficient condition for local invertibility holds. A central result of the observability theory establishes that the Jacobian matrix *J* (Φ *_s_*) can be interpreted as the *observability matrix*, *O_s_*, defined for nonlinear systems [31]. This definition for *O_s_* provides a clear link between the observability of a dynamical system, from an observable *s*, and the existence of singularities in Φ *_s_*, which seemed to be lacking in the literature. In the context of nonlinear systems, there are regions in phase space that are naturally less observable than others.

Following [31], the degree of observability attained from a given variable is quantified with the respective observability index using a value average along an orbit

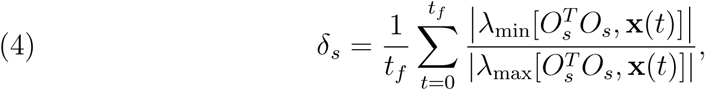

where *t_f_* is the final time considered (without loss of generality the initial time was set to be *t* = 0) and *T* represents the transposition of matrices. The term 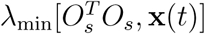 indicates the minimum eigenvalue of matrix 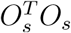 estimated at a point **x**(*t*) (likewise for 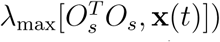. Hence, 0 ≤ *δ_s_* ≤ 1 and the lower bound of *δ_s_*(**x**) is reached when the system is unobservable at point **x**(*t*). It is important to emphasize that the observability indices are local quantities, interpreted as relative values. The stablished average is particularly useful in order to portray an overall picture of the coupling complexity between the dynamical variables.

Each time series arises from a given set of parameters. In this sense, being a function of a dynamical state **x**(*t*), the observability indices are considered local quantities in terms of the parameter values. Given the orbit **x**(*t*), the observability value results from a time average over that orbit. In this sense, the observability indices are considered averaged values along an orbit.

Our present application of the outlined formalism, where the observability matrix is interpreted as the Jacobian matrix of the coordinate transformation in study, *O_s_* = *J* (Φ*_s_*), leads to the computation of the observability indices of the populations *C* = *x*_1_, *T* = *x*_2_ and *I* = *x*_3_. In particular, for *r*_1_ = 0.1842 and *k*_1_ = 0.0001, the observability indices averaged over a trajectory are *δ_x_*_1_ = 0.000634602…, *δ_x_*_2_ = 0.0000124925…, and *δ_x_*_3_ = 0.000000107349…

From the previous values, the original variables can be ranked in descending degree of observability according to

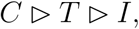

where ▷ means ‘*provides better observability index of the underlying dynamics than*’. As illustrated in Fig. 2, the previous ordering of the observability indices holds for all the populations analyzed in this article. More details about the behavior of the highest observability index, *δ_C_*, in the parameter space are given in Fig. 3.

**FIGURE 2.**
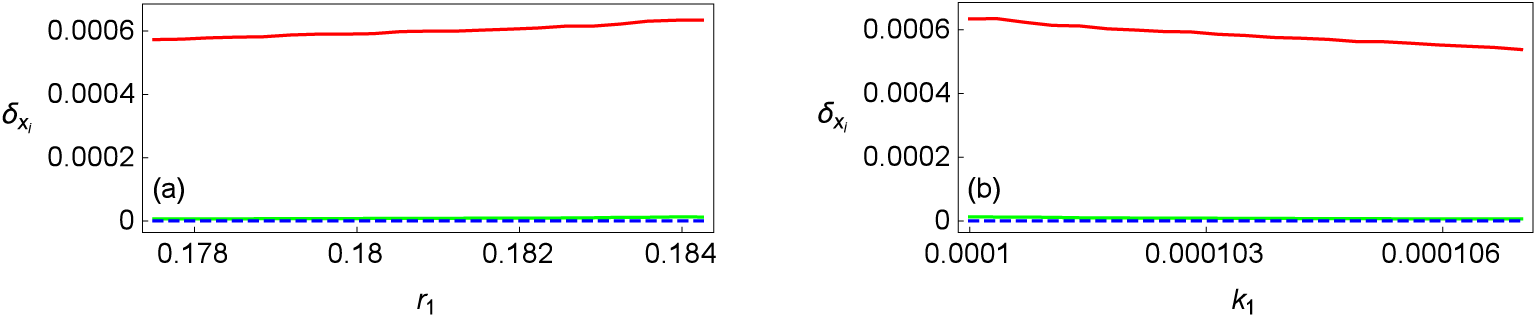
Variation of the three observability indices, *δ_xi_* (*i* = 1, 2, 3), with *C* = *x*_1_, *T* = *x*_2_ and *I* = *x*_3_, where *δ_C_* > *δ_T_* > *δ_I_*.(*a*) *k*_1_ = 0.0001 and 0.1775 ≤ *r*_1_ ≤ 0.18425; (b) *r*_1_ = 0.1842 and 0.0001 ≤ *k*_1_ ≤ 0.000107.

**FIGURE 3.**
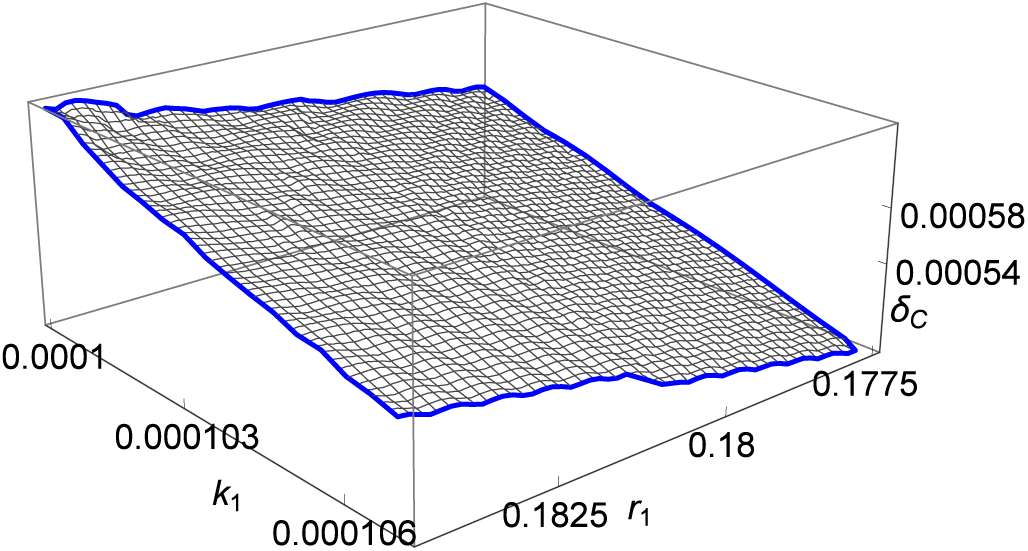
Variation of the highest observability index, *δ_C_* considering 0.1775 ≤ *r*_1_ ≤ 0.18425 and 0.0001 ≤ *k*_1_ ≤ 0.000107.

The three induced phase portraits from the system using the derivative coordinates are displayed in Fig. 4. The computation of the observability indices indicates that variable *C* is the best observable, while *I* is the poorest. The important message of this analysis is that the dynamics of the three-variable HIV-1 model is observed with higher reliability from the population of cancer cells (variable *C*), rather than from the populations of healthy and infected cells (variables *T* and *I*, respectively). The population of healthy cells *T* provides an observability of the dynamics that is less than the one provided by the population of cancer cells *C*, but populations of infected cells *I* is associated with a clearly poor observability (*δ_I_* is smaller than *δ_C_* by three orders of magnitude).

**FIGURE 4.**
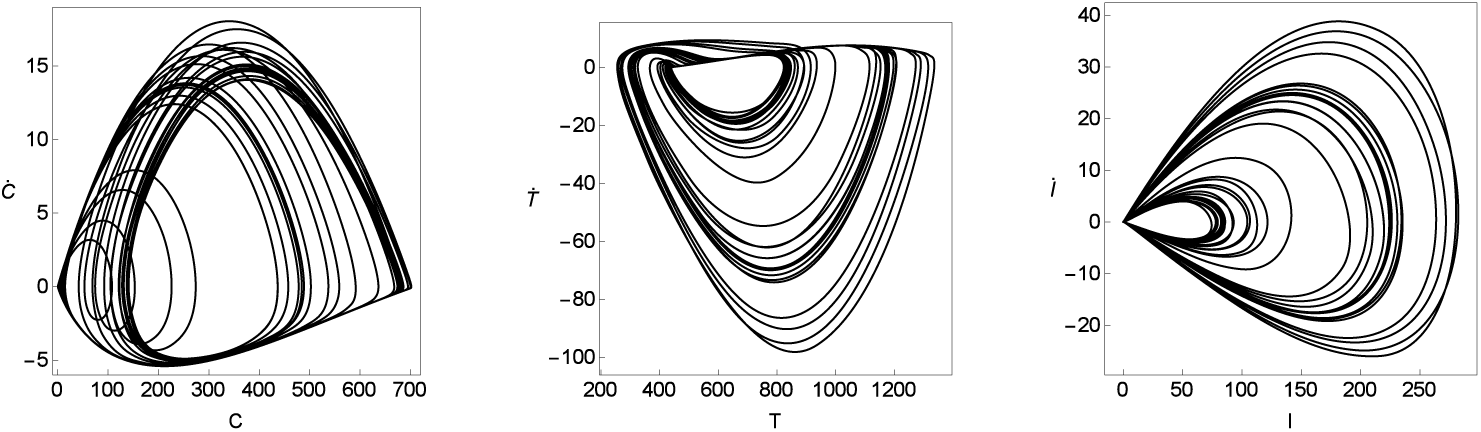
Projections of the dynamics on the (*C*, *C*), (*T*,*T*), and (*I*, *I*) planes used to compute observability indices. Here, we use *r*_1_ = 0.1842 and *k*_1_ = 0.0001.

Our previous result indicates that for systems with HIV-1 infected cells and cancer cells, the variable with largest observability is the population of cancer cells. This should be the population monitored in possible experiments tracking cells populations *in vivo* in order to have the better observable for further analyses. In this sense, some techniques have been developed to monitor different cell populations *in vivo* in mice [33]. The availability of these techniques together with humanized mouse models for HIV-1 [34] could provide a good framework for obtaining time series of cancer cells under these interactions. Such time series could be then used to characterize the underlying dynamics by means of the so-called attractor reconstruction techniques (see Discussion Section).

A 3D-reconstructed attractor would be the result of the representation of the points (*X*_1_,*X*_2_,*X*_3_), with coordinates given by the transformation Φ*_s_*, (result not shown). Only this 3D-representation can be directly compared with the original 3D attractor. The 2D-representations of (*X*_1_,*X*_2_) are different entities, they are just phase portraits, and not necessarily similar to the 3D attractor. In the observability theory, the dynamical states (*X*_1_,*X*_2_) are used to provide the first brief glances over the complexity of the orbits.

In the next paragraphs, we devote a special attention to the dynamical variable *C*. In order to gain insights about the long time behavior of variable *C*, we display in Fig. 5 bifurcation diagrams as a result of the variation of the control parameters *r*_1_ and *k*_1_. Overlapped to the bifurcations diagrams, we display the maximum Lyapunov exponent computed numerically following a standard method [35].

**FIGURE 5.**
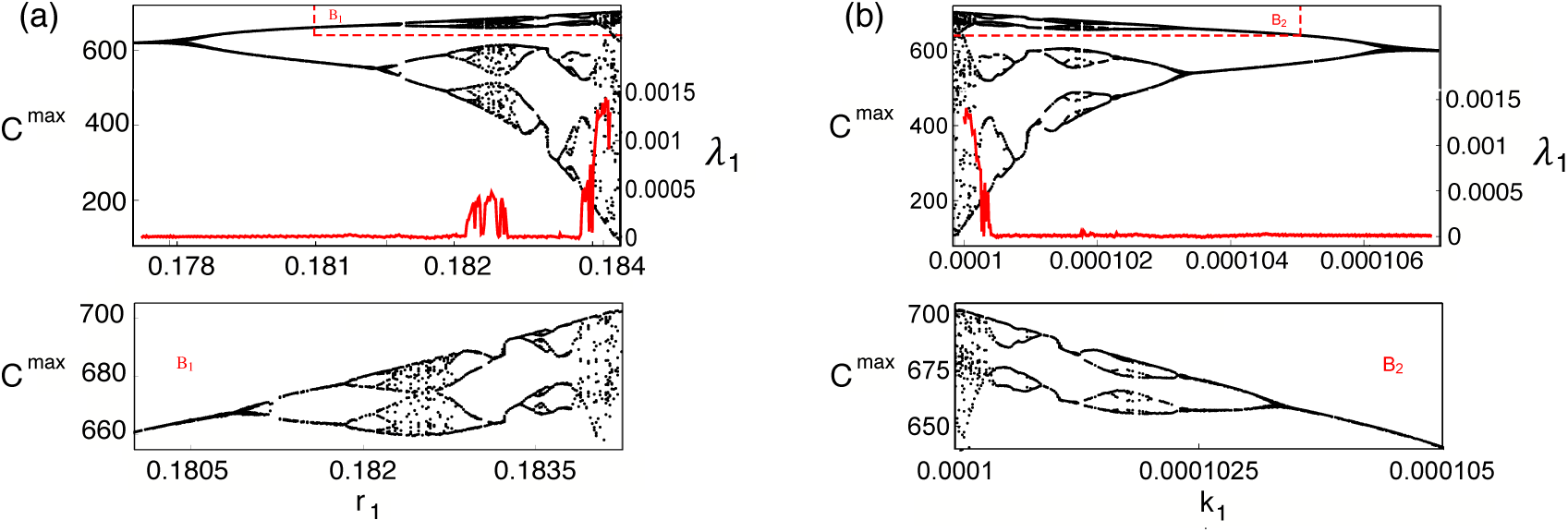
Bifurcation diagrams obtained from the successive *C*^max^. In (a) we tune parameter *r*_1_ within the range 0.1775 ≤ *r*_1_ ≤ 0.18425 using *k*_1_ = 0.0001. In (b) we set *r*_1_ = 0.1842 tuning *k*_1_ in the range 0.0001 ≤ *k*_1_ ≤ 0.000107. The maximum Lyapunov exponent, λ_1_, is shown (in red) overlapped on the bifurcation diagrams. Enlarged views of the boxes *B*_1_ and *B*_2_ are displayed in the lower panels.

## 5. The homotopy analysis methodology and the analytic solutions

For the sake of clarity, we outline in this section a brief description of the HAM (please see [15, 18, 21] and references therein). The analytical approach will be used in a sequence of intervals, giving rise to the step homotopy analysis method. In the context of HAM, each equation of a system of ordinary differential equations

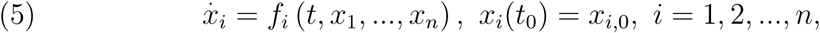

can then be written in the form

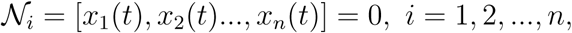

where 𝒩_1_, 𝒩_2_, …, 𝒩*_n_* are nonlinear operators, *x*_1_(*t*), *x*_2_(*t*)…, *x_n_*(*t*) are unknown functions and *t* denotes the independent variable. The analytical procedure starts with a construction of the so-called *zero^th^-order deformation equation*

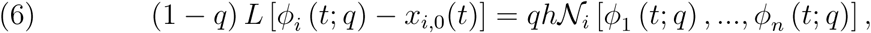

where *q* ∈ [0, 1] is called *the homotopy embedding parameter*, *h is the convergence control parameter*, *L* is an auxiliary linear operator, *x_i_*, _0_ (*t*) are initial guesses for the solutions and *ϕ_i_* (*t*; *q*) are unknown functions. It is clear that when *q* = 0 and *q* = 1, it holds

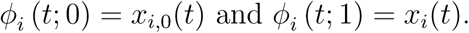

Following (6), when q increases from 0 to 1, the function *ϕ_i_* (*t*; *q*) varies from the initial guess *x_i_*,_0 (*t*)_ to the solution *x_i_*(*t*). Expanding *ϕ_i_* (*t*; *q*) in MacLaurin series with respect to *q* at *q* = 0, we get

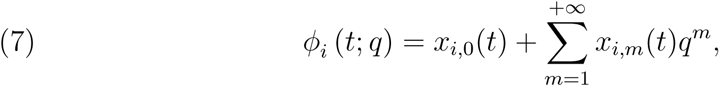

where the series coefficients *x_i_* are defined by

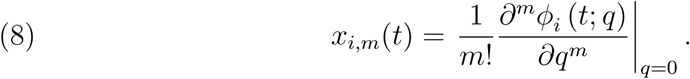

Considering the convergence of the homotopy series (7) and using the relation *x*_i_(*t*) = *φ_i_* (*t*; 1) we obtain the so-called homotopy series solutions

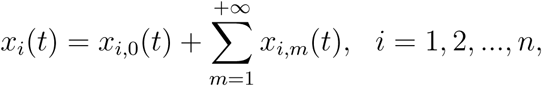

which are precisely the solutions of the original nonlinear equations. Differentiating the *zero^th^*-order deformation Eqs. (6) m times with respect to the homotopy parameter *q*, we obtain the *m^th^*-order deformation equations

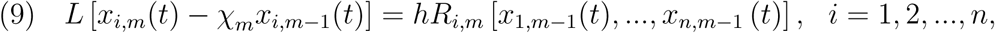

where

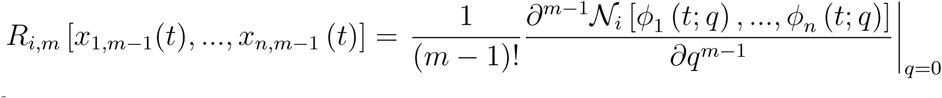

and

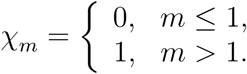

A one-parameter family of explicit series solutions is obtained by solving the linear equations (9). In the presence of some strongly nonlinear problems, it is usually appropriate to apply the HAM in a sequence of subintervals giving rise to the so-called Step Homotopy Analysis Method (SHAM).

### 5.1. Explicit series solution

Following the previous considerations, we are able now to perform an analytical approach of the HIV-1 model by using SHAM. Our goal is to obtain the explicit series solution for *C*, *T*, I and focus our analysis on the analytic solution of the dynamical variable *C*, which represents the population of cancer cells.

Let us consider Eqs. (1)–(3) subject to the initial conditions *C*(0) = *IC*_1_, *T*(0) = *IC*_2_, *I*(0) = *IC*_3_, which are taken in the form *C*_0_(*t*) = *IC*_1_, T_0_(t) = IC_2_, I_0_(t) = *IC*_3_, as our initial approximations of *C* (*t*), *T* (*t*) and *I* (*t*), respectively. In all of our analyses, we will consider *IC*_1_ = 678, *IC*_2_ = 452, *IC*_3_ = 0.25.

As auxiliary linear operators, we choose

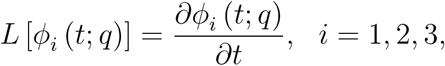

with the property *L* [*C_i_*] = 0, where *C_i_* (*i* = 1, 2, 3) are integral constants. The equations of the HIV-1 model lead to the following nonlinear operators 𝒩_1_, 𝒩_2_ and *N*_3_

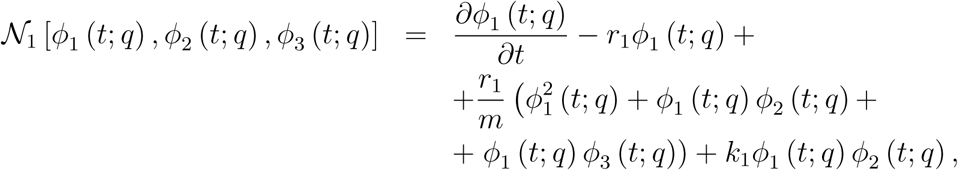

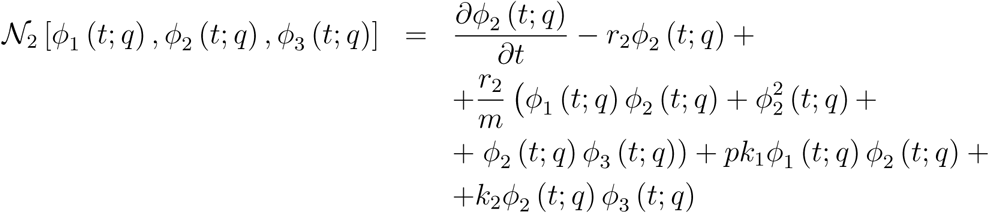

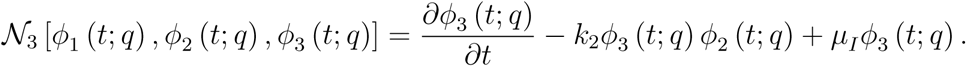

Considering *q* ∈ [0,1] and h the non-zero auxiliary parameter, the *zero^th^*-order deformation equations are

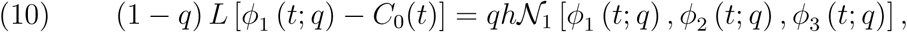

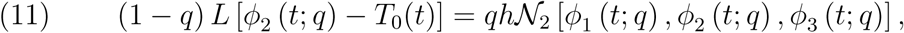

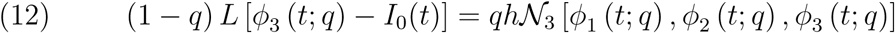

and subject to the initial conditions

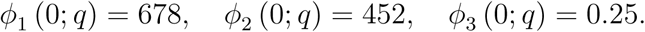

Obviously, for *q* = 0 and *q* =1, the above *zero^th^*-order equations (10)-(12) have the solutions

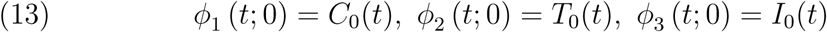

and

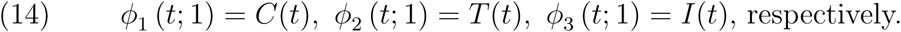

When *q* increases from 0 to 1, the functions *φ*_1_ (*t*; *q*), *φ*_2_ (*t*; *q*) and *φ*_3_ (*t*; *q*) vary from *C*_0_(*t*), *T*_0_(*t*) and *I*_0_(*t*) to *C* (*t*), *T* (*t*) and *I* (*t*). As a result of expanding *φ*_1_ (*t*; *q*), *φ*_2_ (*t*; *q*) and *φ*_3_ (*t*; *q*) in MacLaurin series with respect to *q*, we obtain the homotopy series

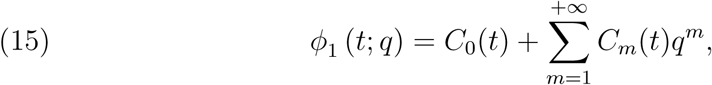

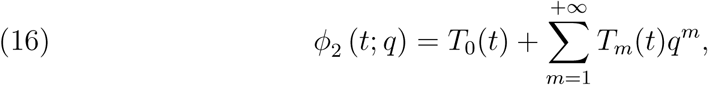

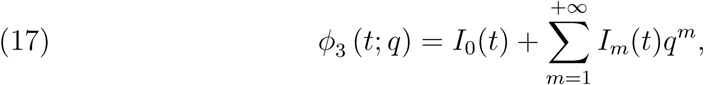

in which

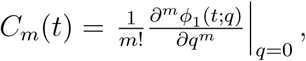

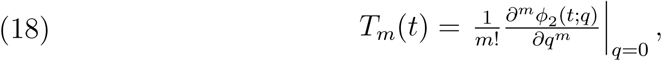

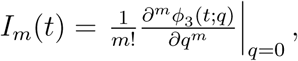

and where *h* is chosen in such a way that these series are convergent at *q* = 1.

Therefore, considering Eqs. (13)-(18), we end up obtaining the homotopy series solutions

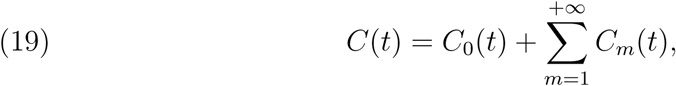

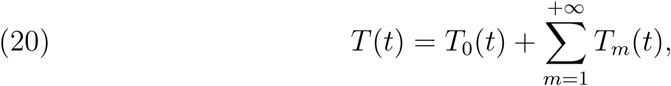

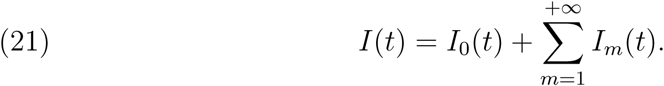

Differentiating the *zero^th^*-order Eqs. (10)-(12) m times and using the properties, where *D_m_* is the *m^th^*-order derivative in order to the homotopy parameter *q*,

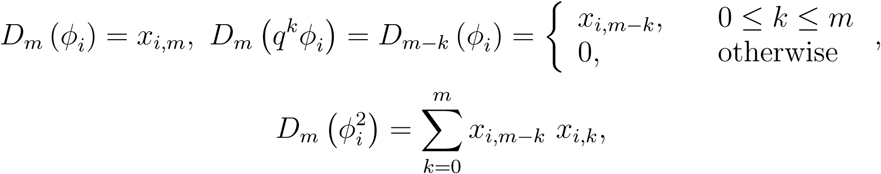

and

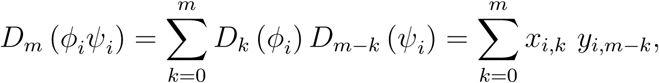

we obtain the *m^th^*-order deformation equations

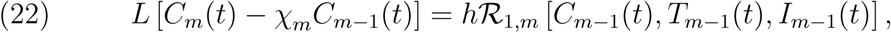

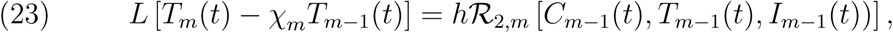

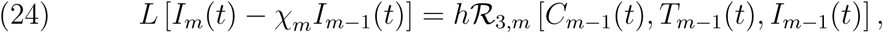

with the following initial conditions

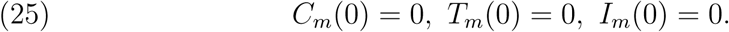

Defining the vector 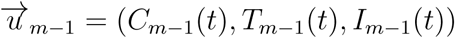 we derive

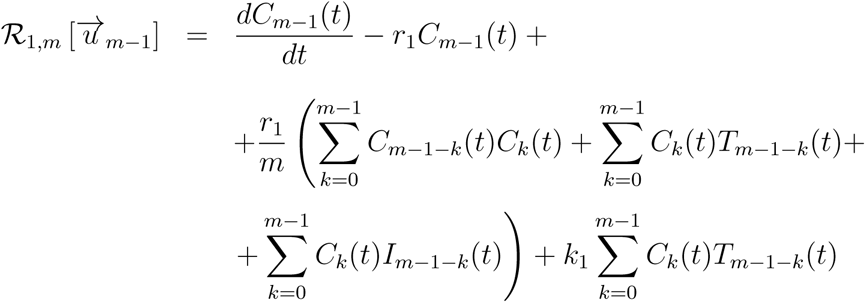

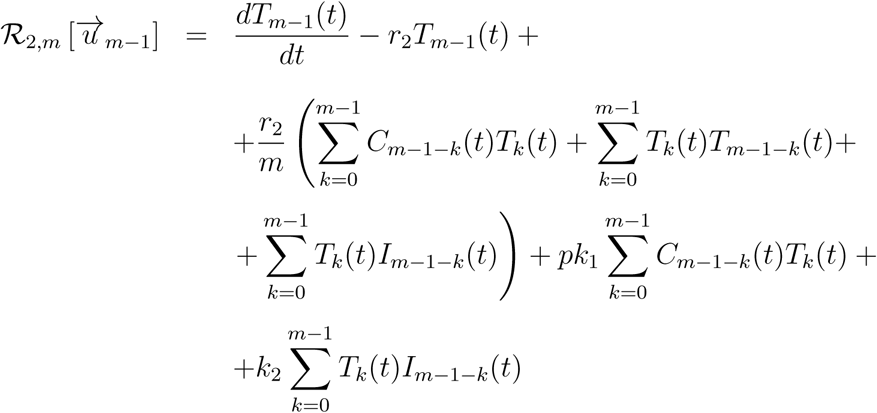

and

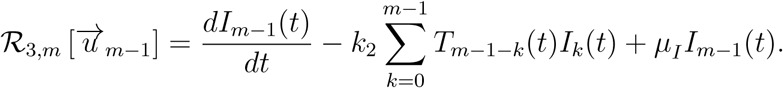

According to the notations and definitions provided above, the solution of the linear *m^th^*-order deformation equations (22)-(24) at initial conditions (25), for all m > 1, becomes

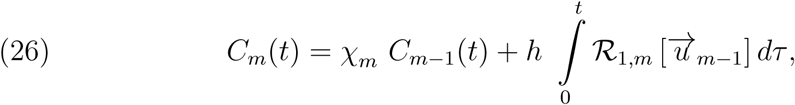

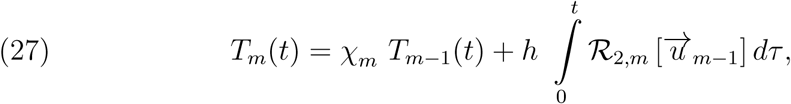

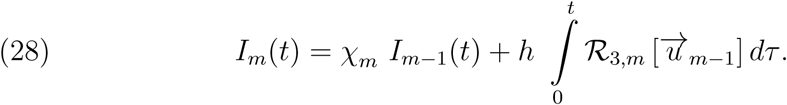

As an example, we present some initial terms of the series solutions (corresponding to *m* = 1 and *m* = 2)

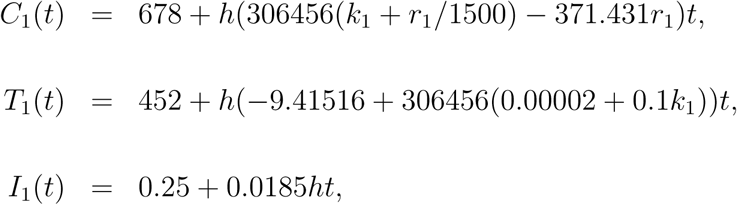

and

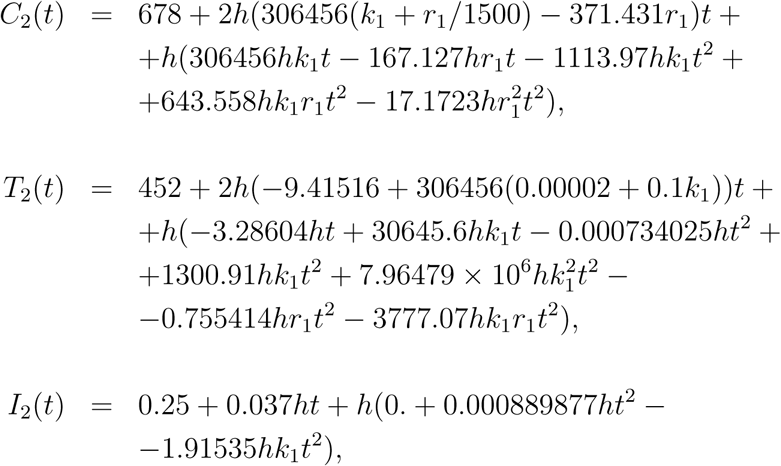

where *h* is the convergence control parameter, *r*_1_ is the uncontrolled proliferation rate of the cancer cells and *k*_1_ is the immune system’s killing rate of cancer cells. At this moment, it is easy to obtain terms for other values of *m*. In particular, truncating the homotopy series (19)-(21) we get the *M*^th^-order approximate analytic solution (which corresponds to a series solution with *M* +1 terms)

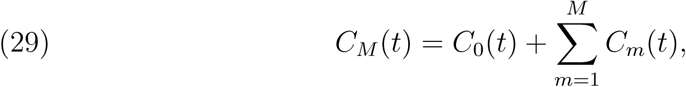

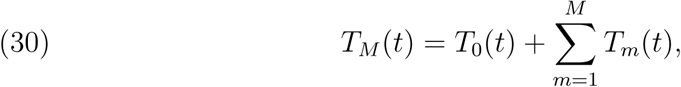

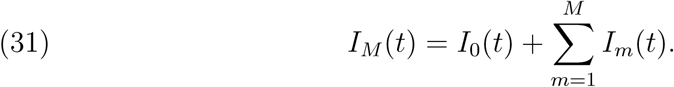

The *exact solutions* are given by the limits

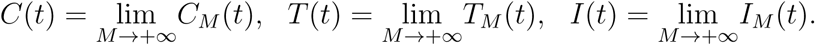

Within the purpose of having an effective analytical approach of Eqs. (1)-(3) for higher values of *t*, we use the step homotopy analysis method, in a sequence of subintervals of time step Δ*t* and the 9-term HAM series solutions (8*^th^*-order approximations)

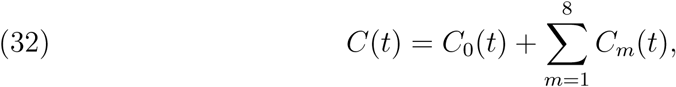

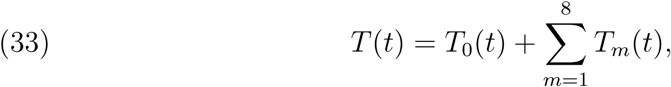

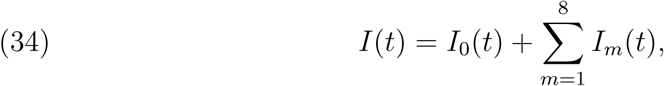

at each subinterval. Accordingly to SHAM, the initial values *C*_0_, *T*_0_ and *I*_0_ change at each subinterval, i.e., 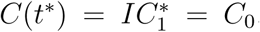, 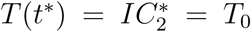 and 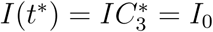 and the initial conditions should be satisfied for all *m* ≥ 1. Therefore, the terms *C_m_*, *T_m_* and *I_m_*, exhibited before as an example for *m* = 1, 2, take now the form

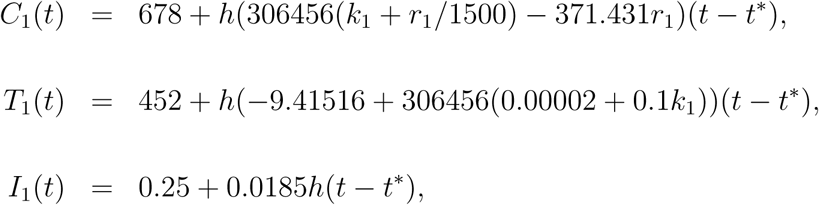

and

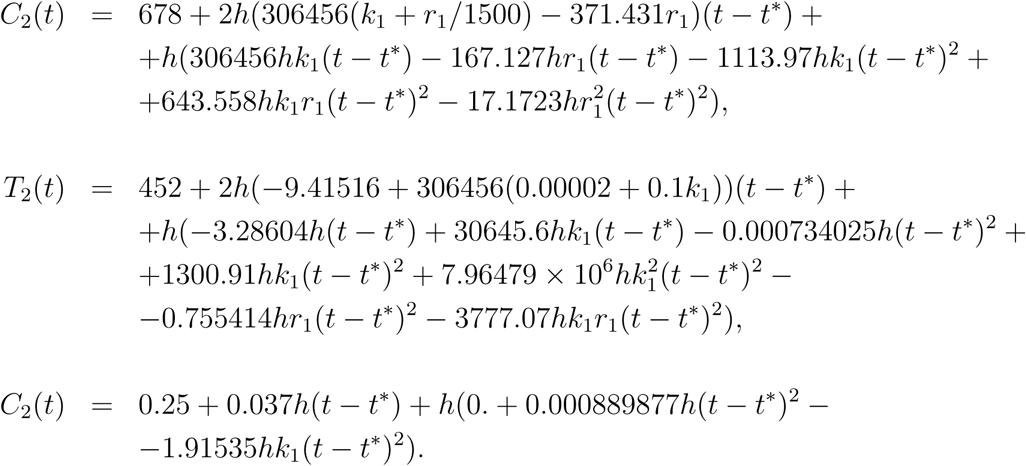

In a similar way, identical changes occur for the other terms. As a consequence, the analytical approximate solution for each dynamical variable is given by

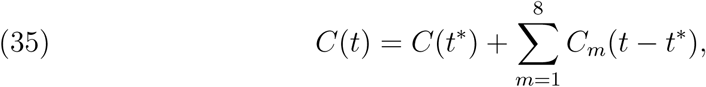

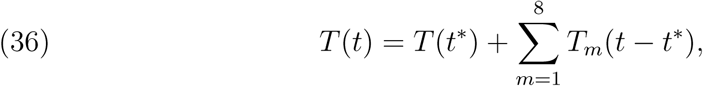

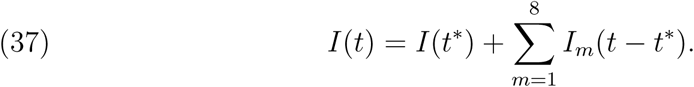

In general, we only have information about the values of *C*(*t*), *T*(*t*) and *I*(*t*) at *t* = 0, but we can obtain the values of *C*(*t*), *T*(*t*) and *I*(*t*) at *t* = *t*\* by assuming that the new initial conditions are given by the solutions in the previous interval. Another illustration of the use of SHAM can be seen in [36].

The homotopy terms depend on both the physical variable *t* and the convergence control parameter *h*. The artificial parameter *h* can be freely chosen to adjust and control the interval of convergence, and even more, to increse the convergence at a reasonable rate, fortunately at the quickest rate. This concept plays a key role in the HAM and is generally used to gain sufficiently accurate approximations with the smallest number of homotopy terms in the homotopy series (29)-(31). In fact, the use of such an auxiliary parameter clearly distinguishes the HAM from other perturbation-like analytical techniques.

How to find a proper convergence control parameter h to get a convergent series solution or, even better, to get a faster convergent one? In the following section, an optimal homotopy analysis approach is decribed to improve the computational efficiency of the homotopy analysis method for nonlinear problems.

### 5.2. An optimal homotopy analysis approach of solutions

Using an optimal approach, the homotopy analysis method might be applied to solve complicated differential equations with strong nonlinearity. Firstly, with the purpose of determining an interval of convergence and the optimum value of *h*, corresponding to each dynamical variable, we state in *Subsection 5.2.1* a recent convergence criterion addressed in [37]. Finally, in *Subsection 5.2.2* an exact Squared Residual Error (SRE) is defined and efficiently used to find optimal convergence values for the convergence control parameter *h*.

It is found that all optimal homotopy analysis approaches greatly accelerate the convergence of series solution.

### 5.2.1. Interval of convergence and optimal value from an appropriate ratio

Let us consider *k* + 1 homotopy terms *x*_0_(*t*), *x*_1_(*t*), *x*_2_(*t*), …, *x_k_*(*t*) of an homotopy series

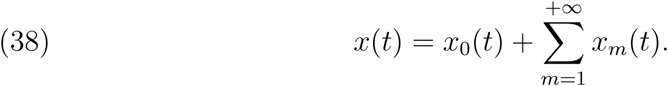

For a preassigned value of parameter *h*, the convergence of the homotopy series is not affected by a finite number of terms. Therefore, it is sufficent to keep track of magnitudes of the ratio defined by

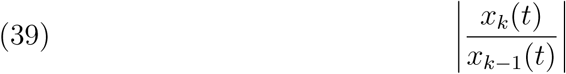

and whether it remains less than unity for increasing values of *k*. Taking (39), and requiring this ratio to be as close to zero as possible, we can determine an optimal value for the convergence control parameter *h*. For such a value, the rate of convergence of the homotopy series (38) will be the fastest (and as a consequence, the remainder of the series will rapidly decay). For a prescribed h, if the ratio is less than unity, then the convergence of HAM is guaranteed. In other words, this is a sufficient condition for the convergence of the homotopy analysis method. This implies that in the cases where the limit for the ratio in (39) cannot be reached or tends to unity, the method may still converge or fail to do so. It is appropriate to search for an optimum value of *h*, i.e., a value of *h* that gives rise to a ratio (39) as small as possible. Taking a time interval Ω, the ratio

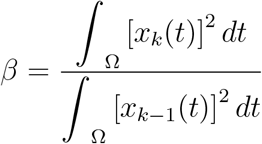

represents a more convenient way of evaluating the convergence control parameter *h*. In fact, given an order of approximation, the curves of ratio *β* versus *h* indicate not only the effective region for the convergence control parameter *h*, but also the optimal value of *h* that corresponds to the minimum of *β*. Now, plotting *β* versus *h*, as well as by solving

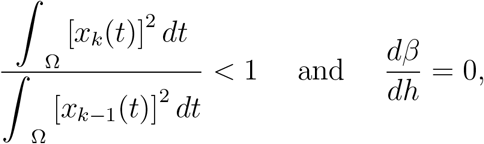

the interval of convergence and the optimum value for parameter *h* can be simultaneously achieved.

**Table 1.** Intervals of convergence of *h* and the respective optimum values *h*,* corresponding to the dynamical regimes presented in Fig.6 (for *r*_1_ = 0.1842 and *k*_1_ = 0.0001).

| β-Curves | Intervals of convergence and optimal values of $h$ |
| --- | --- |
| $\beta_C$ | $-1.55274 < h_C < -1.09974$<br>$h_C^* = -1.18485$ |
| $\beta_T$ | $-1.66625 < h_T < 0$<br>$h_T^* = -1.28944$ |
| $\beta_I$ | $-0.190914 < h_I < 0$<br>$h_I^* = -0.122441$ |

As an illustration at the order of approximation *M* = 8, the curves of ratio *β* versus *h*, corresponding to *C*(*t*), *T*(*t*) and *I*(*t*) *(β_C_* vs *h_C_*, *β*_T_ vs *h_T_* and *β_I_* vs *h_I_*, respectively), are displayed in Fig. 6. In Table 1, we exhibit the intervals of convergence of *h* and the respective optimum values *h*\* corresponding to the dynamical regime presented in Fig. 6.

**FIGURE 6.**
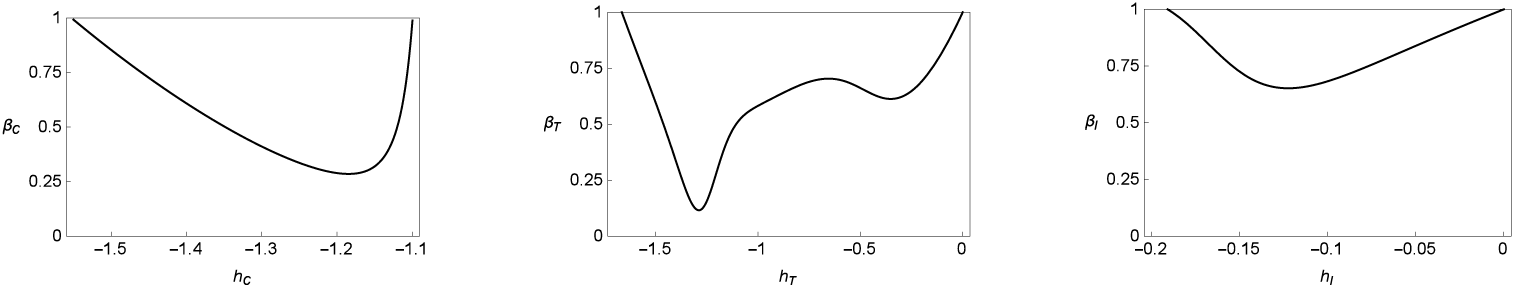
(From left to right) The curves of ratios *β_C_*, *β_T_* and *β_I_* versus *h_C_*, *h_T_* and *h_I_*, respectively, corresponding to a 8*^th^*-order approximation of solutions *C*(*t*), *T*(*t*) and *I*(*t*) for *r*_1_ = 0.1842 and *k*_1_ = 0.0001. The optimum value of *h*, *h*\*, gives rise to the minimum value of *β*.

Indeed, the use of such ratio, by solving the inequality mentioned above, allows us to obtain the exact interval of convergence for the artifitial parameter *h* and, in addition, it yields an optimal value. This represents a central advantage in the study of the convergence of HAM. In Fig. 7 we show the comparison of the SHAM analytical solutions for *C*, *T* and *I* with the numerical results using precisely the optimum values presented in Table 1.

**Figure 7.**
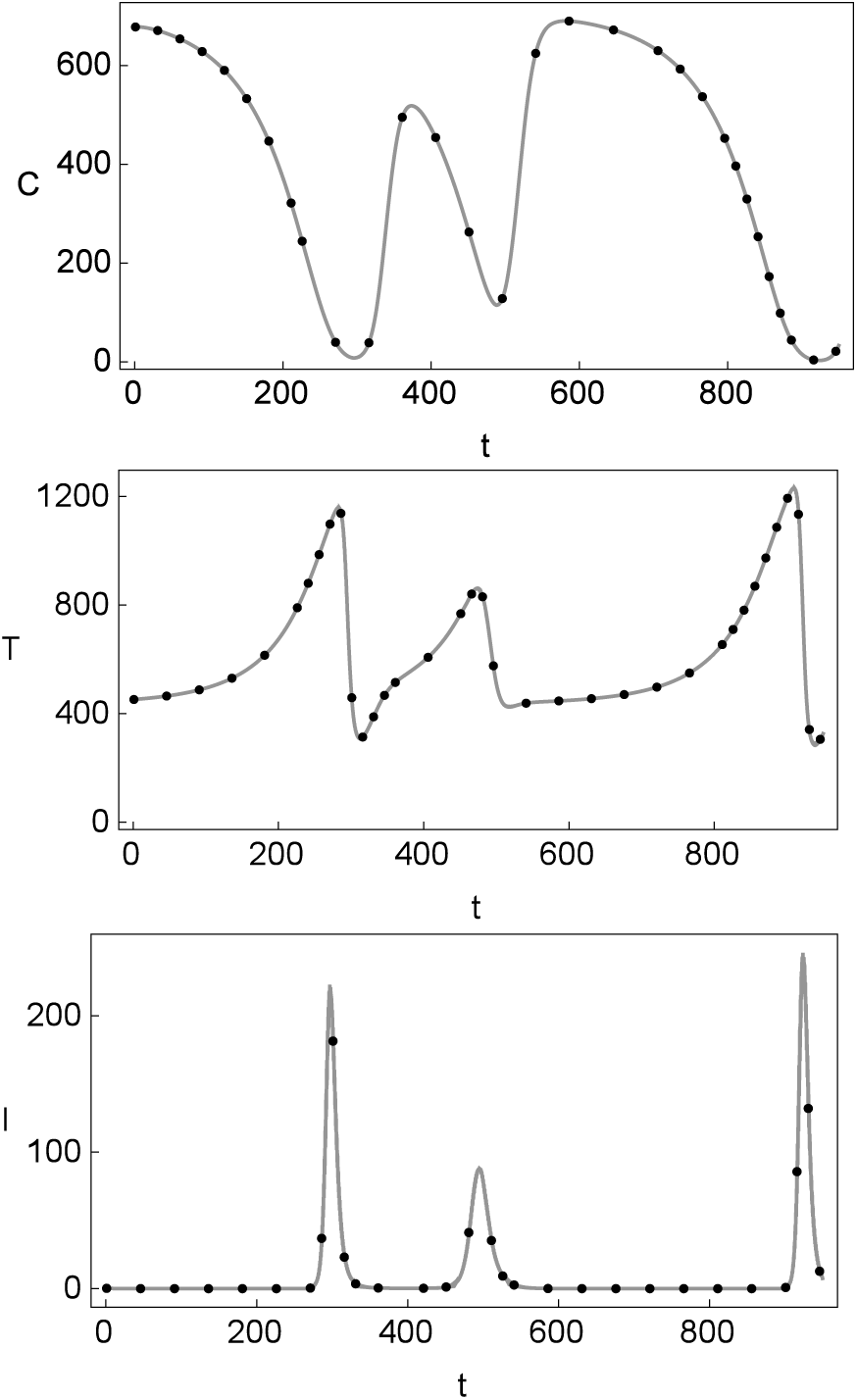
Comparison of the SHAM analytical solutions (35)-(37) of *C*, *T* and *I* (solid lines) with the respective numerical solutions (dotted lines) of the HIV-1 cancer model. The value of the control parameters are *r*_1_ = 0.1842 and *k*_1_ = 0.0001.

### 5.2.2. Squared residual error and different orders of approximation

A procedure to check the convergence of a homotopy-series solution is to substitute this series into the original governing equations and initial conditions, and then to evaluate the corresponding squared residual errors - the more quickly the residual error decays to zero, the faster the homotopy-series converges. In this context, an error analysis is performed in the following lines.

Taking the expressions (29)-(31), let us consider *φ_C_* (*t*,*h_C_*) = *C_M_*(*t*), *φ*_T_ (*t*, *h_T_*) = *T_M_*(*t*), *φ_I_* (*t, h_I_*) = *I_M_*(*t*). With the substitution of these solutions into Eqs. (1)-(3), we are able to construct *Residual Error* (RE) functions as follows:

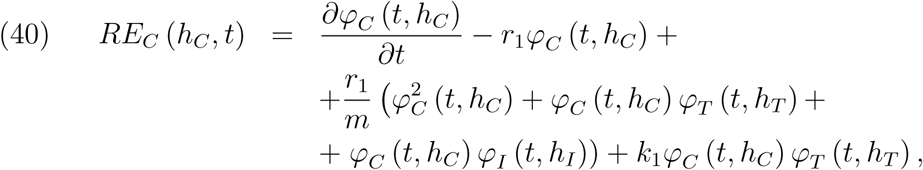

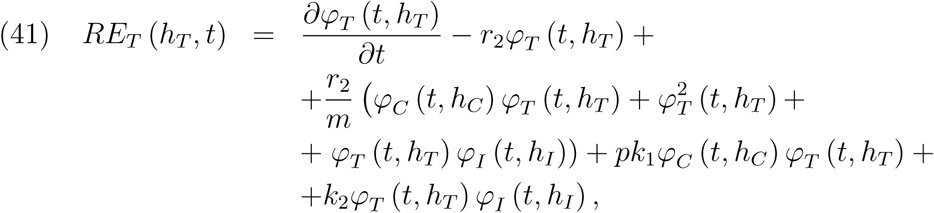

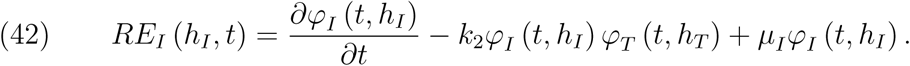

In 2007, Yabushita *et al.* [38] suggested an *optimization method* for convergence control parameters. Their work is based on the Squared Residual Error. Inspired by this approach, and following the studies carried out in [39, 40], we consider the exact *Squared Residual Error* (SRE) for the *M^th^*-order approximations to be

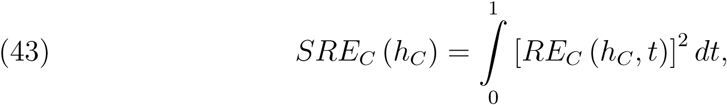

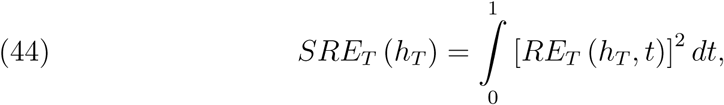

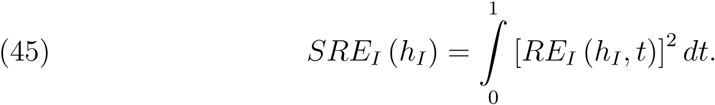

Values of *h_C_*, *h_T_* and *h_I_* for which *SRE_T_* (*h_T_*), *SRE_I_* (*h_T_*) and *SRE_I_* (*h_I_*) are minimum can be obtained. For a given *M*^th^-order of approximation, the optimal value of *h_C_*, *h_T_* and *h_I_* are given by the nonlinear algebraic equations

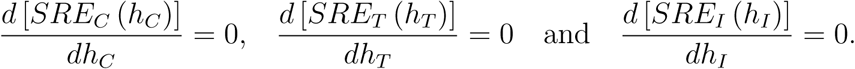

The optimal values for all of these considered cases are 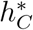, 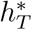 and 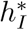. The curves of *SRE_C_*, *SRE_T_* and *SRE_I_* regarding different orders of approximation, namely *M* = 6, *M* = 8 and *M* = 10, are show in Fig. 8. Central information regarding the orders of approximation, optimal values of *h*_C_, *h*_T_, *h_I_* and minima of the respective squared residual error functions is summarized in Table 2.

**FIGURE 8.**
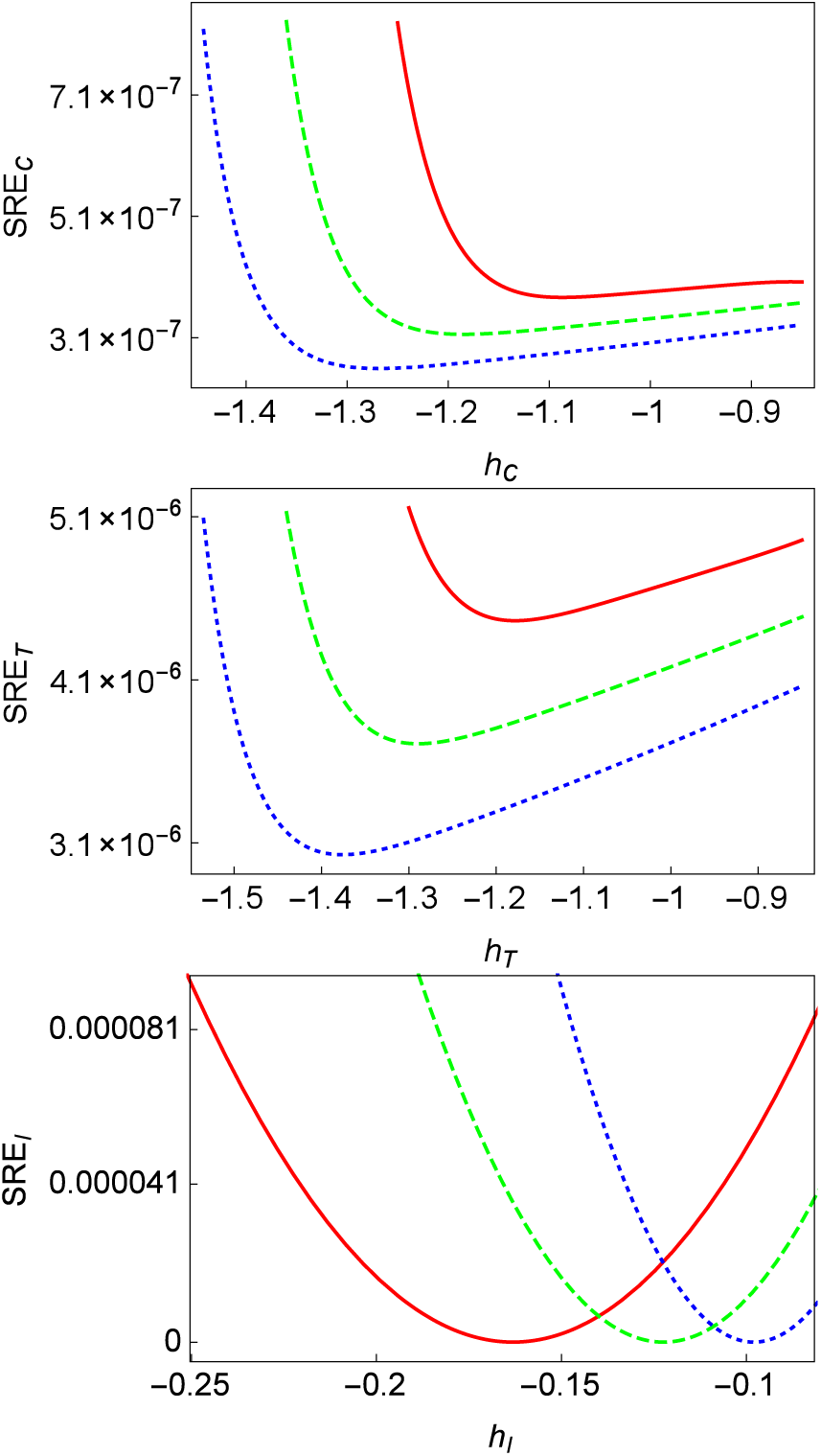
Exact Squared Residual Error functions, *SRE_C_*, *SRE_T_* and *SRE_I_*, versus *h_c_*, *h_T_* and *h_I_*, respectively. These functions correspond to different orders of approximation for the solutions *C*(*t*), *T*(*t*) and *I*(*t*). Solid red line: 6*^th^*-order opproximation; Dashed green line: 8*^th^*-order opproximation; Dotted blue line: 10*^th^*-order opproximation (*r*_1_ = 0.1842 and *k*_1_ = 0.0001). Each optimum value *h*\* gives rise to the minimum value of the SRE.

**Table 2.** Orders of approximation, optimal values of *h_C_*, *h_T_*, *h_I_* and minima of the respective squared residual error functions, corresponding to the dynamical regime presented in Fig. 8 (*r*_1_ = 0.1842 and *k*_1_ = 0.0001).

| $M$, order of approximation of $C(t)$ | Optimal value $h_C^*$ | Minimum value of $SRE_C$ |
| --- | --- | --- |
| 6 | -1.08697 | $3.76497 \times 10^{-7}$ |
| 8 | -1.18485 | $3.15689 \times 10^{-7}$ |
| 10 | -1.26551 | $2.5934 \times 10^{-7}$ |
| $M$, order of approximation of $T(t)$ | Optimal value $h_T^*$ | Minimum value of $SRE_T$ |
| 6 | -1.17838 | $4.46271 \times 10^{-6}$ |
| 8 | -1.28944 | $3.70809 \times 10^{-6}$ |
| 10 | -1.37576 | $3.02756 \times 10^{-6}$ |
| $M$, order of approximation of $I(t)$ | Optimal value $h_I^*$ | Minimum value of $SRE_I$ |
| 6 | -0.163095 | $5.14312 \times 10^{-8}$ |
| 8 | -0.122441 | $4.78534 \times 10^{-8}$ |
| 10 | -0.0979792 | $4.57679 \times 10^{-8}$ |

This analysis provides an illustration of how our understanding of a model arising in the context of biology can be directly enhanced by the use of numerical and analytical techniques, for different combinations of control parameters and time.

## 6. Discussion

In this paper we have provided new insights into the study of an HIV-1 model, which mimics the interactions between populations of cancer cells, healthy *CD*4+ T lymphocytes and infected *CD*4+ T lymphocytes. The rich and complex behavior of this model allowed us to apply different theoretical approaches. The presence of chaos in predator-prey cancer models has been reported in several theoretical articles [41, 42, 32]. The direct detection of chaos in cancer dynamics is hard to seek, since long time series of population changes of specific cells should be obtained *in vivo* or *in vitro*. Techniques to detect and quantify multiple distinct populations of cells circulating simultaneously in the blood of living animals are being currently developed. For instance, an optical system for two-channel, two-photon flow cytometry *in vivo* [33]. The possibility of studying HIV-1 infection in humanized mouse models [34] with tumors using this technique could allow an experimental setup to monitor the population of cancer cells in HIV-infected mice (see below).

Despite the difficulty of experimentally identifying chaos in tumor dynamics, some mathematical models based on the biological interactions between tumor, healthy, and immune cells have revealed the possibility of this behavior using parameters values matching some biological evidences, and could be thus considered as being qualitatively validated with experimental data. The model analyzed in this article considers predator-prey-like interactions between cancer and immune cells, also adding HIV-1 infected immune cells. This model as well, reveals the presence of chaos using some parameters from clinical and mathematical models [13].

After analytically proving the boundedness of the trajectories in the system’s attractor, we have studied the complexity of the coupling between the dynamical variables with the quantification of the observability indices. Our calculations revealed that the highest observable variable is given by the population of cancer cells. Hence, cancer cells should be the ones monitored in experiments in order to have reliable data for possible attractor reconstruction techniques [32]. For example, the time-delay embedding (i.e., Taken’s embedding theorem) or differential embedding [45]. The time-delay embedding technique has been successfully applied for attractor reconstruction in several systems such as the Belousov-Zhabotinskii reaction [43] and epidemics [44], amog others. Once a chaotic attractor is reconstructed, several dynamical and topological measures (like Lyapunov exponents or fractal dimension) could be performed. Such an approach could be used to characterize the attractor governing coexistence among cancer cells, *CD*4+ T cells, and HIV-infected T cells. As mentioned above, scalar time series of cancer cell populations could be obtained *in vivo* using the two-channel, two-photon technique in model organisms.

We have identified different dynamical behaviors of the system varying two biologically meaningful parameters: *r*_1_, representing the uncontrolled proliferation rate of cancer cells and *k*_1_ denoting the immune system’s killing rate of cancer cells. The changes in parameter *r*_1_ reveal regions with periodic and chaotic behavior, which have been identified with the computation of the maximum Lyapunov exponent.

Nonlinear equations are significantly more difficult to solve than linear ones, especially in terms of analytical methods. In general, there are two standards for a *satisfactory* approach of nonlinear equations: (i) it can always give approximation expressions *efficiently*; (ii) it can guarantee that approximation expressions are *accurate* enough in the studied region of biophysical parameters. Using these two standards as a criterion, we have successfully applied the homotopy analysis method (HAM) to construct the explicit series solution of the HIV-1 model incorporating AIDS-related cancer cells. The HAM solution contains the auxiliary parameter h, which gives a simple way to adjust and control the convergence region of the resulting series solution. In order to increase the computational efficiency, an optimal homotopy analysis approach was developed to obtain optimal values for the convergence-control parameter *h* by means of an appropriate ratio and the definition of an exact Squared Residual Error. This analysis provided a fast convergence of the homotopy series solution and illustrated that the homotopy analysis method indeed satisfies the two standard aspects, (i) and (ii), mentioned previously.

The results presented in this article are likely to inspire applications of the HAM analytical procedure for solving highly nonlinear problems in theoretical biology. This study provides another illustration of how an integrated approach, involving numerical evidences and theoretical reasoning within the theory of dynamical systems, can directly enhance our understanding of biologically motivated models.

## Acknowledgements

This work was partially funded by FCT/Portugal through UID/MAT/04459/2013, and by “la caixa” Foundation (JS).

